# Rapid Alkalinization Factor (RALF) gene family genomic structure and transcriptional regulation during host-pathogen crosstalk in *Fragaria vesca* and *Fragaria* x *ananassa* strawberry

**DOI:** 10.1101/858928

**Authors:** Francesca Negrini, Kevin O’Grady, Marko Hyvönen, Kevin M. Folta, Elena Baraldi

**Affiliations:** Laboratory of Plant Pathology and Biotechnology, DISTAL-University of Bologna, Bologna Italy; Departmnent of Horticultural Science, University of Florida, Gainesville, Florida, United States of America; Department of Biochemistry, University of Cambridge, Cambridge, United Kingdom

## Abstract

Rapid Alkalinization Factor (RALF) are cysteins-rich peptides ubiquitous in plant kingdom. They play multiple roles as hormone signals, starting from root elongation, cell growth, pollen tube development and fertilization. Their involvement in host-pathogen crosstalk as negative regulator of immunity in *Arabidopsis* has also been recognized. In addition, RALF peptides are secreted by different fungal pathogens as effectors during early stages of infections. Campbell and Turner previously identified nine RALF genes in *F. vesca* v1 genome. Here, based on the recent release of *Fragaria x ananassa* genome and *F. vesca* reannotation, we aimed to characterize the genomic organization of the RALF gene family in both type of strawberry species according to tissue specific expression and homology with *Arabidopsis*. We reveal the presence of 13 RALF genes in *F. vesca* and 50 in *Fragaria x ananassa,* showing a non-homogenous localization of genes among the different *Fragaria x ananassa* subgenomes associated with their different TE element contents and genome remodeling during evolution. *Fragaria x ananassa* RALF genes expression inducibility upon infection with *C. acutatum* or *B. cinerea* was assessed and showed that, among fruit expressed *RALF* genes, *FaRALF3-1* was the only one upregulated after fungal infection. *In silico* analysis and motif frequency analysis of the putative regulatory elements upstream of the *FaRALF3* gene was carried out in order to identify distinct pathogen inducible elements. Agroinfiltration of strawberry fruit with 5’ deletion constructs of the *FaRALF3-1* promoter identified a region required for *FaRALF3* expression in fruit, but did not identify a region responsible for fungal induced expression.

## Introduction

In plants, several small secreted peptides (SSPs) function as hormone signalling molecules to respond to internal and external stimuli [1]. SSPs are known to be involved in different processes, ranging from organs growth to biotic and abiotic responses [2, 3].

Rapid alkalinisation factors (RALFs) are cysteine-rich SSPs originally identified for their ability to rapidly alkalinize tobacco cell culture (4). They are ubiquitous in plant kingdom with 37 members identified in *Arabidopsis thaliana* genome alone [5, 6]. RALF genes are translated as pre-pro-peptides and activated in the apoplast through proteolytic cleavage. Besides the signal sequence necessary for extracellular extrusion, canonical RALF proteins contain distinctive amino acid motifs, such as the RRILA motif for S1P protease recognition [7] and the YISY motif, important for signaling cascade activation [8, 9]. In addition, four conserved cysteines form two disulfide bonds stabilise the mature RALF proteins. Based on these features, RALFs have been classified into four major clades [6]; clades I, II and III contain typical RALF peptides, whereas clade IV groups the most divergent RALF peptides, lacking RRILA and YISY motifs, and in some cases containing only three cysteines.

RALF peptides bind to the *Catharanthus roseus* Receptor Like Kinases 1 - like family protein (CrRLK1L) known to be involved in cell expansion and reproduction throughout the plant kingdom [10]. The large *Cr*RLK1L receptor family which includes FERONIA (FER) receptors, previously reported to interact with *Arabidopsis RALF1* and *RALF23* [11, 12], Buddah’s Paper Seal 1, 2 (BUPS1/2), ANXUR1, 2 (ANX1/2) protein, which interact through their ectodomain and bind to RALF4 and 19 in pollen the tube (13) and THESEUS1 (THE1), the RALF34 receptor in roots (14). Binding RALF peptides to *CrRLK1L* receptors also involve other interacting partners such as Lorelei-like-Glycosylphosphatidylinositol-Anchored proteins (LLG1,2,3) [9, 15] and Leucin-Rich Repeat Extensins (LRX) have been reported to bind RALF4/19 in the pollen tube [16, 17] and to interact with FER, as part of cell wall sensing system responsible for vacuolar expansion and cellular elongation (18). RALF binding to receptors leads to a number of different intracellular signaling events involving different molecular components, mostly still unidentified. However, it is known that in *A. thaliana* binding of RALF1 to FER receptor results in plasma membrane H(+)-adenosine triphosphatase 2 phosphorylation, causing the inhibition of protein transport and subsequent apoplastic alkalinization (11).

RALF peptides regulate a variety of different functions such as cell expansion (11), root growth [19], root hair differentiation [20, 21, 22] stress response [23], pollen tube elongation and fertilization [13, 16, 24, 25]. Recently, it was reported that in *A. thaliana,* RALF peptides act as negative regulators of the plant immune response to bacterial infection [12], since the binding of processed RALF23 to the FER receptor inhibits the formation of the complex between the immune receptor kinases EF-TU RECEPTOR (EFR) and FLAGELLIN-SENSING 2 (FLS2) with their co-receptor BRASSINOSTEROID INSENSITIVE 1–ASSOCIATED KINASE 1 (BAK1), which is necessary to initiate immune signalling. Interestingly, biologically-active RALF homologs have also been identified in fungal plant pathogens, possibly following interspecies horizontal gene transfer, pointing to a role for fungal RALF genes as virulence factors [26]. In fact, a RALF-homolog fungal peptide is fundamental for host alkalinisation and infection by *Fusarium oxysporum* [27]. Since for many pathogenic fungi alkalinisation is important for activation of virulence factors and successfully infection of plant tissues, secreted RALF peptides may act as initial effectors to promote host alkalinisation at early stage infection, when hyphal biomass is not sufficient to secrete a large quantities of ammonia [28]

Strawberry (*Fragaria x ananassa*) is an important crop but is highly susceptible to different fungal and bacterial pathogens severely affecting crop yield [29]. In particular post-harvest molds are difficult to manage since, even if they originate from field infection, they are acquired during flowering stage but can manifest after a long asymptomatic quiescent period on ripe fruits. Severe strawberry fruit post-harvest molds are caused by *Colletotrichum acutatum,* causal agent of anthracnose disease and *Botrytis cinerea,* causal agents of grey mold.

Expression analysis of *Fragaria x ananassa* fruits infected with *C. acutatum* at two different ripening stages revealed an increase in the expression of a RALF gene in the susceptible fruit at early stage infection [30]. Overexpression of the *Fragaria x ananassa* RALF gene homolog of *A. thaliana RALF33* through transient agroinfiltration increases strawberry fruit susceptibility to anthracnose, causing a major fungal biomass growth on fruits and the induction of the host immune response [30, 31]. Upregulation of RALF genes during plant infection has also been observed in mature red tomato fruits (*Solanum lycopersicum)* infection by *Colletotrichum gleosporioides* and in rice upon *Magnaporthe oryzae* infection [32]. This suggests a role for RALF gene expression as a susceptibility factor in fungal infection. [33].

Furthermore Dobón *et al.* [34], studying the expression pattern of four *Arabidopsis* transcription factors mutants (*at1g66810, pap2, bhlh99, zpf2*) characterized by the increased susceptibility to *B. cinerea* and *Plectosphaerella cucumerina*, observed an upregulation of *RALF23, RALF24, RALF32* and *RALF33* genes. The woodland strawberry RALF gene family members have been previously characterized based on the *F. vesca* genome annotation v1 [6] and nine *FvRALF* genes were identified and grouped in the four RALF clades.

Considering the recent release of a new annotated version (v4.0.a2) of the *Fragaria vesca* genome [35] and of the *Fragaria x ananassa cv. Camarosa* genome sequence v1.0.a1 [36, 37], we set out to explore both newly released genome annotations in order to update the RALF gene family composition and genomic organization in woodland strawberry (*F. Vesca*), and the four subgenomes of the *Fragaria x ananassa* octoploid strawberry. For this, genome localization, phylogenetic and transcriptional analysis were conducted, based on available genomic and transcriptomic sources, with the aim to gaining insights into the different functional roles of RALF genes. Furthermore, the inducibility of RALF genes expression upon *Fragaria x ananassa* fruits infection with *C. acutatum* and *B. cinerea* was studied. *In silico* analysis of the *FaRALF3-1* promoter was conducted also in order to identify putative pathogen responsive motifs. We then tested *in vivo* if progressively truncated *FaRALF3-1* promoter fragments could induce reporter genes expression in agroinfiltarted strawberry fruits infected with *C. acutatum*.

## Materials and Methods

### RALF family genes identification, phylogenetic tree analysis, clade determination and chromosome map

*F. vesca* RALF genes already reported by Campbell and Turner [6] were implemented through keyword gene search ‘RALF’ GDR (v4.0.a2) [35]. The output was compared and integrated. Nucleotide sequences of the 13 *F. vesca* RALF members were used as query for BLASTx on *Fragaria x ananassa cv. Camarosa* Genome v1.0.a1 proteome [38] to find octoploid RALF homologs. *Fragaria x ananassa* RALF peptide sequences were aligned by MUSCLE [39] and the phylogeny was inferred using the Maximum Likelihood method and JTT matrix-based model [40]. The tree with the highest log-likelihood was chosen. The Initial tree was obtained automatically by applying Neighbor-Join and BioNJ algorithms to a matrix of pairwise distances estimated using a JTT model, and then selecting the topology with superior log-likelihood value. Evolutionary analyses were conducted in MEGA X [41]. To determine new members classification in clades, all the RALF peptide sequences available from the Campbell and Turner annotation plus the updated *F. vesca* genes were aligned and phylogentic tree were designed as mentioned above. *Fragaria x ananassa* genes annotation and position were retrieved from GDR (*Fragaria x ananassa* cv. Camarosa genome v1.0.a1), and progenitor lineage were inferred according to [36]. Chromomap package in R [42] was used to create a *Fa*RALF gene chromosome map.

### Expression profile of RALF family genes in *F. vesca* and qRTPCR in *Fragaria x ananassa*

Heatmap expression profile of RALF family genes in different *F. vesca* tissues were designed using heatmap3 package in R [43] from Transcripts Per Kilobase Million (TPM) values calculated by Li et al. [35].

### Infection of Fragaria x ananassa fruits

*Fragaria x ananassa cv. Alba* plants were grown in the greenhouse at 25°C and 16 hours light. White (21 days after anthesis) and red fruits (28 days after anthesis) were harvested and infected or not according to the experimental procedures. Each treatment contained at least three biological replicates. *Colletotrichum acutatum* (Isolate Maya-3, from CRIOF-UniBo fungi collection) and *Botrytis cinerea* strain *B05.10* were grown on PDA plates for 15 days. Detached fruits were dipped for 30 s in a 10^6^ conidia per mL suspension or in water as negative control and incubated for 24 hours at room temperature in plastic bags

### RNA extraction and qRT-PCR expression analysis

The surface of experimental fruits were excised with a scalpel and immediately frozen in liquid nitrogen. RNA was extracted according to Gambino *et al*. [44], run on an 2% Agarose gel and quantified with NanoDrop™ 3300 for respectively integrity and quality control. cDNA was made starting from 1 µg of RNA using Promega ImProm-II™ Reverse Transcription system. qRT-PCR analysis was performed using ThermoFisher MAXIMA SYBR GREEN/ROX QPCR 2x supermix. RALF gene relative expression was calculated using standard curve method and *Elongation Factor* gene as reference (*XM_004307362.2)* Primers for RALF genes expression analysis were designed on *Fragaria x ananassa, F. vesca* subgenome genes and specificity were checked observing a single peak in the dissociation curve for each primer pair. All the primers used for RALFs and transient transformation reporter genes expression analysis are listed in Table S2.

### Statistical Analyses

The numerical values quantifying expression of RALF genes and eGFP reporter, were calculated as average of three independent biological replicates each formed by a group of at least three treated fruits. For eGFP quantification, all mean normalized expression values were expressed relatively to the negative empty pKGWFS7 vector infiltrated fruits. To determine statistical significant samples, Student t-test between samples and relative control was performed.

### *In silico* analysis of predicted regulatory elements in *FaRALF3-1* promoter

For pathogen-induced predicted regulatory elements analysis, pathogen-upregulated genes were retrieved from transcriptional datasets of red strawberry (*Fragaria x ananassa)* fruits after 24 hours post infection with *C. acutatum* [30] and *B. cinerea* [45]. A total of 87 predicted promoters for *C. acutatum* and 97 for *B. cinerea* were analyzed. For each gene, 1500 bp upstream the ATG start codon were analyzed from the *F.vesca* genome v4.0.a1 assembly. MotifLab software v1.08 [46] was used for *in silico* analysis using PLACE database for Motif Scanning [47] and AlignACE algorithm for Motif Discovery [48]. Briefly, Motif Scanning were performed using Simple Scanner program with default parameters. Statistically significant known regulatory elements were calculated performing the same Motif scanning program on randomly generated DNA sequences starting from input predicted promoter sequences using a third order background model. The frequency measured for each cis-acting element on random-DNA was used as background occurrence for statistical significant evaluation using a binomial test with p-value threshold of 0.05. Motif Discovery was performed using AlignACE method with default parameters and motif significance were calculated as mentioned above for Motif scanning method.

### *Fragaria x ananassa* RALF3 promoter characterization

Preliminary study on putative promoter allelic variants were conducted on *Fragaria x ananassa cv. Alba* and *cv. Florida Elyana.* Genomic DNA was extracted from leaves using Wizard® Genomic Purification kit (Promega). Plant Tissue Protocol (Manufacturer protocol 3.E.) was modified adding two consecutive chloroform:isoamyl alcohol (24:1) purification steps after Protein Precipitation solution addition and before 2-propanol precipitation. *FaRALF3-1* putative promoter was amplified using primers For 5’-TGCATCTGTTACATCATCCCTTG-3’ and Rev 5’-GTAGTCGACTCTCCCATCTTG-3’, cloned into pGEM®-T easy vector (Promega). Five clones for each variety were sequenced and aligned with *Fragaria x ananassa cv. Camarosa* genomic sequence available from GDR, using Clustal Omega.

### Progressive truncated promoter cloning and *Agrobacterium*-mediated transient transformation

*FaRALF3-1* upstream sequence was PCR-amplified starting from *Fragaria x ananassa cv. Elyana* genomic DNA, using primers 5’-GGGGACCACTTTGTACAAGAAAGCTGGGTNCTGAAAGGACAAAACATTTTCT-3’ as reverse primers for all promoter fragments, 5’-GGGGACAAGTTTGTACAAAAAAGC AGGCTNNTGCATCTGTTACATCATCCCTTG-3’ as forward for whole promoter fragment (T6), 5’-GGGGACAAGTTTGTACAAAAAAGCAGGCTNNTGCTTAAGTGGCTCTCAAAG-3’ as forward for 400 bp fragment (T4) and 5’-GGGGACAAGTTTGTACAAAAAAGCAGGCTNNCCGCTAAGTGGTTCAATTCA-3’ as forward for 200bp fragment (T2). Truncated FvRALF3 promoter constructs, and double tandem p35S promoter as positive control, were cloned into pDONR222 using Gateway BP reaction and consequently cloned into pKGWFS7 (S1 Figure) vector by LR Reaction. Obtained vectors were then introduced into chemically competent *Agrobacterium tumefaciens* strain EHA105 by heat shock transformation. Briefly, after been frozen in liquid nitrogen, cells were thawed for 5 min at 37°C, 1 µg of plasmid DNA were added, then cell were incubated at 30°C for 2 hours in agitation and plated. Positive colonies were then grown in selective media (Rinfampycin 100 µg/mL and Spectinomycin 50 µg/mL) until culture reached an OD_600_ of 0.8. Cells were then collected through centrifugation and pellet was resuspended in fresh MMS medium (Murashige and Skoog Basal Medium 4.4 g/L plus sucrose 20g/l) until an OD_600_ of 2.4 was reached, at the end acetosyringone (4’-Hydroxy-3’,5’-dimethoxyacetophenone) at the final concentration of 200 µM was added to the culture. At least three white attached fruits, for each condition, were agroinfilitrated using a needle syringe until *Agrobacterium* culture filled strawberry fruit tissues. Five days after agroinfiltration, fruit were harvested and infected with *C. acutatum* conidial suspension or mock-inoculated with water, according to experimental procedure, as was described above. After 24 hours post infection fruits were dissected and one half was used for RNA extraction and eGFP expression analysis, and the other half was used for histochemical assay of GUS activity.

### Histochemical GUS assay

Surface tissue and longitudinal sections of infected and mock-infected fruits were cut with a razor blade and dipped in GUS staining solution (50 mM Na-phosphate [pH 7.5], 10 mM EDTA, 1 mM 5-bromo-4chloro-3-indolyl-glucuronide (X-gluc), 0.1% Triton X-100, 0.5 mM potassium ferricianide and 5% (w/v) polyvinylpyrrolidone-40 (PVP)). Strawberry tissues were incubated overnight at 37 °C, and then kept at 4 °C in absolute ethanol until being photographed.

### 3D modelling of FaRALFs interaction with MRLK and LLG2 proteins

Homology models of FaRALF3, in complex with FERONIA and LLG2 were generated using Modeller [49](v9.19) package using the complex of *A. thaliana* RALF23, LLG2 and FERONIA (PDB: 6a5e) as the template. ClustalX was used to create alignments of different components of the complex: FaRALF3 (GDR: snap_masked-Fvb2-2-processed-gene-47.50-mRNA-1), *Fragaria x ananassa* FERONIA MRLK47 (GDR: Uniprot: A0A1J0F5V4) and *Fragaria x ananassa* LLG2 (GDR: maker-Fvb3-4-snap-gene-34.65-mRNA-1) with the *A. thaliana* proteins in the *A. thaliana* complex. PyMOL Molecular Graphic Systems version 1.2r3ore (Schröedinger, LLC) was used for the analysis of the homology models and the figures.

## Results and Discussion

### Identification of RALF gene family members in Fragaria vesca

RALF peptides belonging to 51 plant species have been previously classified in four clades depending on the sequence similarity [6]. Typical distinctive amino acid sequence motifs, such as the RRILA proteolytic cleavage site [7] and the YISY receptor binding site, are present in RALF peptides of clade I to III, and missing in clade IV, which contains more divergent peptides. Nine RALF genes have been previously identified in the *Fragaria vesca* v1 genome.

In order to identify new members of RALF gene family in *Fragaria vesca*, the recent genome annotation (v4.0.a2) [35] was searched using ‘RALF’ as keyword gene name, revealing the presence of 13 RALF genes. The identified *F. vesca* RALF (*FvRALF*) genes are named with progressive numbers according to their chromosome position, from Chr1 to 6 (Table 1), with one RALF gene in Chr1, two in Chr2, −3, and −5, and three in Chr4, and −6. No RALF genes are found in Chr7. Out of the nine RALF genes previously reported by Campbell and Turner [6], eight genes are confirmed both for identity and chromosome position. These are the gene08146 (corresponding to *FvRALF2*), *gene10567 (FvRALF3)*, *gene02376 (FvRALF4)*], *gene02377 (FvRALF5), gene06579 (FvRALF6), gene06890 (FvRALF8), gene10483 (FvRALF9), gene22211 (FvRALF13)* (Table 1). The *gene00145*, previously annotated as gene encoding for a peptide with the typical RALF motifs RRILA and YISY [6], was discharged since in the new v4.0.a2 annotation its sequence corresponds to gene *FvH4_6g07633* encoding for a shorter peptide lacking most of RALF conserved motifs.

**Table 1.** List of RALF genes identified in *F. vesca*.

| RALF genes | v4.0.a2 | Former annotation | Chromosome localization | clade | Arabidopsis homology |
| --- | --- | --- | --- | --- | --- |
| **FvRALF1** | FvH4_1g16140 | gene23829 | Fvb1:9232850..9233915 | IV | *RALF34* |
| **FvRALF2** | FvH4_2g13590 | gene08146* | Fvb2:11869901..11870362 | III | *RALF4* |
| **FvRALF3** | FvH4_2g25351 | gene10567* | Fvb2:20606722..20607381 | II | *RALF33* |
| **FvRALF4** | FvH4_3g09010 | gene02376* | Fvb3:5266873..5268430 | III | *RALF19* |
| **FvRALF5** | FvH4_3g09020 | gene02377* | Fvb3:5269449..5270363 | III | *RALF19* |
| **FvRALF6** | FvH4_4g13190 | gene06579* | Fvb4:16794712..16795349 | II | *RALF32* |
| **FvRALF7** | FvH4_4g13250 | gene06566 | Fvb4:16854377..16855101 | III | *RALF24* |
| **FvRALF8** | FvH4_4g18001 | gene06890* | Fvb4:21972673..21974193 | II | *RALF33* |
| **FvRALF9** | FvH4_5g21290 | gene10483* | Fvb5:12816493..12816858 | II | *RALF33* |
| **FvRALF10** | FvH4_5g26840 | gene41610 | Fvb5:18214922..18215453 | IV | *RALF25* |
| **FvRALF11** | FvH4_6g36290 | gene42389 | Fvb6:28540907..28541125 | IV | *RALF32* |
| **FvRALF12** | FvH4_6g41850 | gene42489 | Fvb6:32780751..32781347 | IV | *RALF5* |
| **FvRALF13** | FvH4_6g06520 | gene22211* | Fvb6:3818620..3820001 | I | *RALF33* |

V4.0.a2 annotation according to Li et al.,[35]; Former annotation corresponds to v1.0.a1 style annotation and asterisk (\*) indicate genes previously identified in v1 *F. vesca* genome [6] and confirmed in the v4.0.a2 annotation version; clade classification according to Campell and Turner [6].

The *FvRALF* genes members were aligned using Clustal Omega (S2 Figure) and classified in the four clades according to Campbell and Turner [6] (Table 1). Overall, one gene (*FvRALF13*) was included in clade I, four in clade II (*FvRALF3, FvRALF6, FvRALF8, FvRALF9*), four in clade III (*FvRALF2, FvRALF4,FvRALF5, FvRALF7*) and four in clade IV (*FvRALF1, FvRALF10, FvRALF11, FvRALF12*).

The *FvRALF* genes include two members homologous to *Arabidopsis AtRALF32* (*FvRALF11, FvRALF6)*, two homologous to *AtRALF19* (*FvRALF4, FvRALF5)*, four homologous to *AtRALF33* (*FvRALF3, FvRALF8, FvRALF13 and FvRALF9)*, and one respectively to *AtRALF4* (*FvRALF2)*, *AtRALF5* (*FvRALF12), AtRALF24* (*FvRALF7), AtRALF25* (*FvRALF10)* and *AtRALF34* (*FvRALF1)*. All *FvRALF* genes are predicted to contain a single exon and no introns. Interestingly, the gene *FvRALF3* is transcribed from a putative Natural Antisense Transcript (NAT) generating region and its complementary sequence encodes for the 3’ untranslated region of a heat shock factor binding protein gene (*FvH4_2g25350)*.

### Identification, evolution and chromosome organization of RALF genes family member in Fragaria x ananassa

*F. vesca* RALF gene sequences were used as query sequences against a database of predicted proteins (v1.0.a1 Proteins source [6]) in *Fragaria x ananassa cv. Camarosa* (*Fxa*). Fifty RALF members were identified in the *Fxa* octoploid strawberry (S1 Table). Paralogous genes (orthologs to particular *FvRALF* genes) were identified in the *Fxa* subgenomes from alignment (S3 Figure) and phylogenetic analysis (Fig 1A) and progenitors lineage were inferred from chromosome localization (Fig 1B) according to Edger *et al.* [36]. Fourteen genes out of these 50 are localized in the *F. vesca* subgenome, 15 members in the *F. nipponica* subgenome, 13 in the *F. iinumae* subgenome and only eight in the *F. viridis* subgenome (Fig 1C).

**Fig 1:**
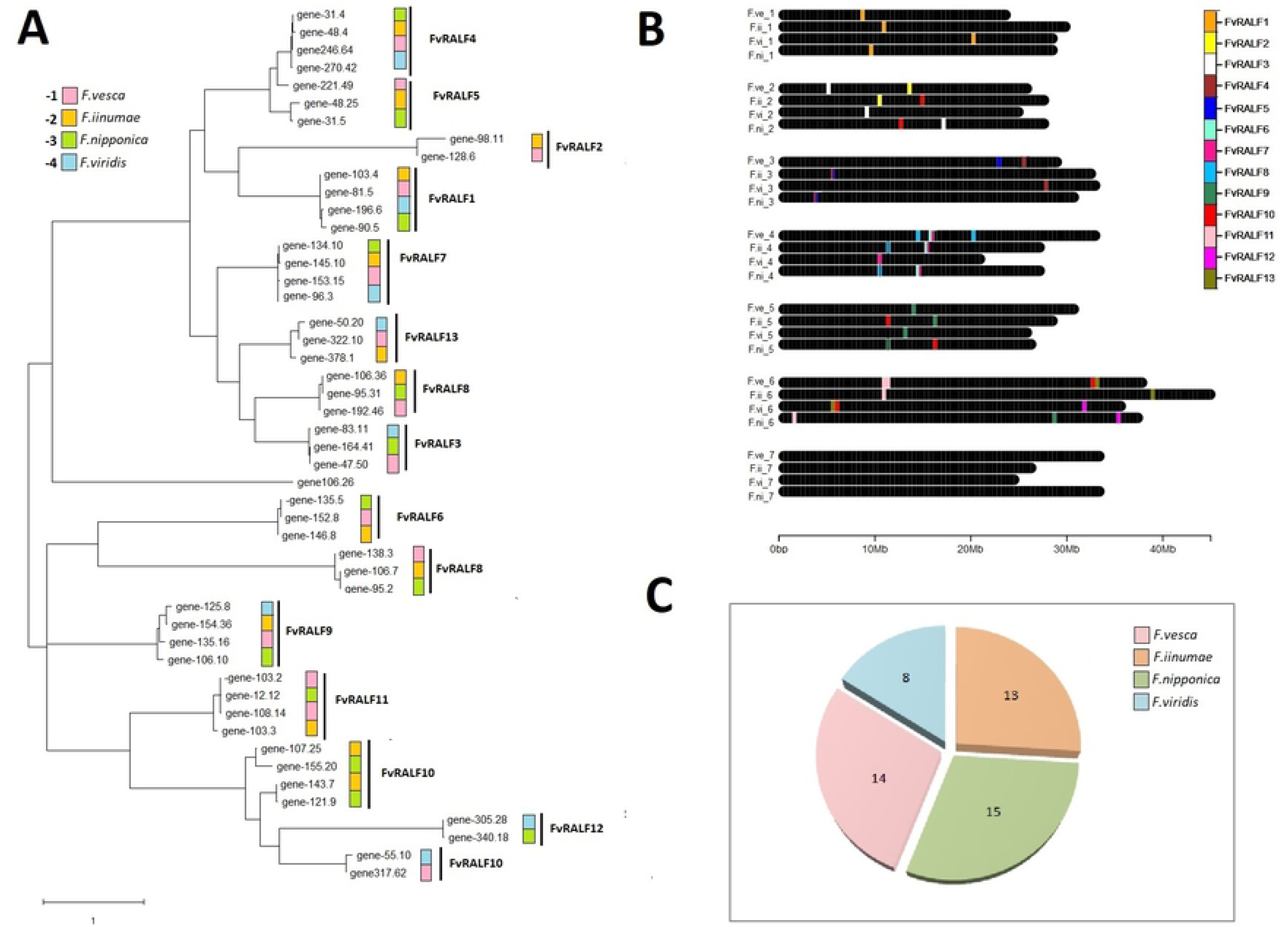
*Fragaria x ananassa (Fxa)* RALF genes phylogentic analysis, evolution and chromosome organization. (a) Phylogenetic tree was built aligning 50 *Fa*RALF peptides sequences using MUSCLE. The tree is drawn to scale, with branch lengths measured in the number of substitutions per amino acidic site in the peptide sequences. Gene annotations refer to *Fragaria x ananassa cv. Camarosa* v1.0.a1 and are listed in Table S1. Progenitor lineage were inferred from gene chromosome localization (b), and pink was used for *F. vesca* subgenome, green for *F. nipponica,* light blue for *F. viridis* and orange for *F. iinumae.* Color legend reports also code number used to name different *FaRALF* genes according to subgenome lineage, as is reported in TableS1. (b) *FaRALF* genes chromosome spatial organization in the octoploid genome. (c) Pie chart showing total FaRALF gene members present in the four subgenomes.

*Fxa RALF* genes *(FaRALF)* were named based on corresponding *FvRALF* orthologs and the subgenome localization, with progressive numbers from 1 to 4 to indicate *F. vesca*, *F. iinumae*, *F. nipponica* and *F. viridis* progenitors, respectively (accordingly to Edger *et al.*[36]), and progressive letters to nominate genes orthologous to the same *FvRALF* genes and localized on the same chromosome (e.g. *FaRALF8-2a* is one of the two *Fxa* genes orthologous to *FvRALF8* mapping on *F. iinumae* subgenome).

Gene homology and chromosome localization analysis showed that only four out of 13 *FvRALF* genes, namely *FvRALF1, FvRALF4, FvRALF7, FvRALF9,* have orthologs in all the four subgenomes. For all the other cases the *FvRALF* orthologs are not represented in all the different *Fxa* subgenomes, probably due to gene loss events occurred during evolution or polyploidy adjustment (Fig 1A). In particular, the *F. viridis* derived subgenome has the lowest number of RALF gene members and is lacking genes orthologous to *FvRALF5, FvRALF8, FvRALF6, FvRALF8, FvRALF11* and *FvRALF2*. On the other hand, similarly to *F. vesca*, no RALF genes are localized in Chr7 of the different progenitors. Moreover in *Fxa* genome some RALF genes probably underwent duplication events. For example *FaRALF11-1a* and *FaRALF11-1b* are both orthologous to *FvRALF11* and are positioned close together on Chr6 (*F.vesca* subgenome). Another atypical gene organization is found for *FaRALF5-1a* and *FaRALF5-1b* genes, located on Chr3 (*F.vesca* subgenome), which are annotated as single genes but contain two tandem RALF conserved domains, suggesting that a duplication event occurred during genome evolution. In addition, the *FaRALF7-4* gene is predicted to encode for a 325 aa protein; containing a conserved RALF domain within the first 104 amino acids and a domain homologous to chloroplastic NADPH-dependent aldehyde-reductase like protein, from aa 124 to 325. This incongruous arrangement of typically unrelated protein domains suggest this gene is a result of shuffling events during evolution. Genes *FaRALF9-3b* and *FaRALF9-3c*, occur as NAT element on Chr 5 (*F. nipponica* subgenome), as was observed in *F.vesca* for *FvRALF3*.

The *F. viridis* derived subgenome contains the least number of RALF genes. This was also the case for R gene family in *Fxa* [36]. However, in contrast to the R gene family, there is not a clear dominance of *F.vesca* progenitor in the *FaRALF* gene family composition, since genes are similarly distributed in the *F. iinumae, F. nipponica* and *F.vesca* subgenomes (Fig1C). As was speculated by Edger *et al*. [36] the lack of RALF genes in *F. viridis* subgenome could be related to the higher TE content of this subgenome which can cause both higher mutation rates and gene loss.

*Fxa* RALF genes classification in clades is consistent with diploid woodland strawberry respect to RALF clade distribution, with 15 genes in clade IV, 17 genes in clade III, 15 in clade II and three in clade I (S1 Table).

### Transcriptional dataset analysis of RALF genes in *F.vesca*

To provide insights into the RALF genes members functions in strawberry, available RNA-seq datasets mapped onto the new genome annotation v4.0.a2 by Li *et al.* [35], were analyzed in different tissues and developmental stages. RALF members were grouped based on similar expression profile and hierarchical clustering resulted in four major RALF expression groups (Fig 2): i) RALF genes specifically expressed in mature male gamete *(FvRALF4, FvRALF5, FVRALF10, FvRALF11)*; ii) a gene expressed only in two anther developmental stages (*FvRALF2)* iii) *FvRALF3* and *FvRALF12* genes mainly expressed in roots and in roots infected with *Phytophthora cactorum* iv) genes mainly expressed in different fruit developmental stages (*FvRALF1, FvRALF6, FvRALF7, FvRALF8, FvRALF9* and *FvRALF13)*.

**Fig 2.**
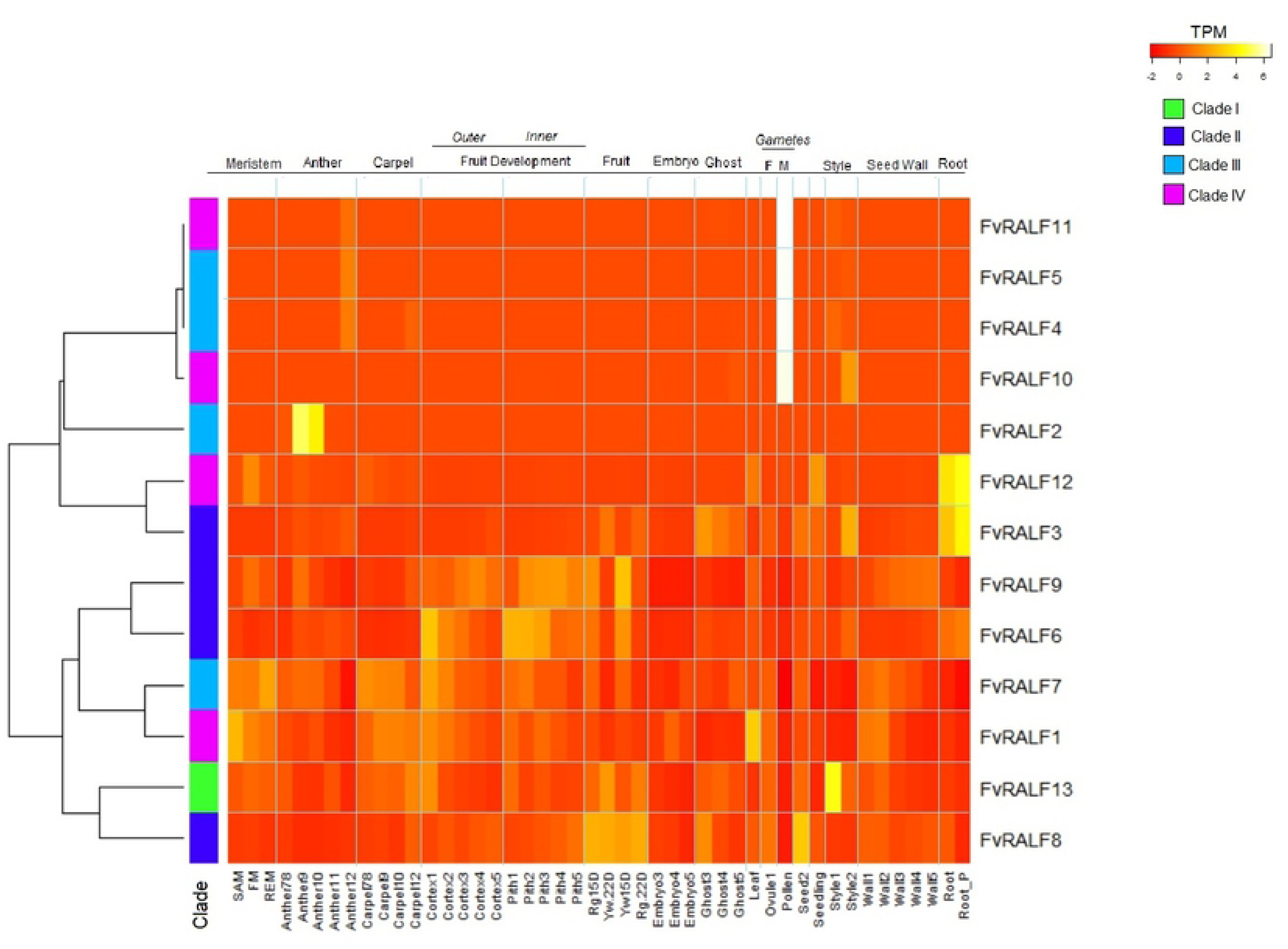
Transcriptional analysis of RALF genes in *F. vesca.* Heatmap of the expression profile measured as Transcription per Kilobase Million (TPM) of *FvRALF* members (rows) in different tissues and developmental stages (column). Labels at the bottom specify the tissues and the stages, and light blue vertical lines divide organs labeled at the top. Dendrogram on the left shows rows relationship according to similar expression profile. RALF members classification in clades are shown: in green clade I, in blue clade II, in light blue clade III and magenta clade IV.

Contrary to what has been observed in *Arabidopsis,* where clade IV RALF genes were highly expressed in flower tissues [6], the woodland strawberry *FvRALF* genes included in each of the four expression groups belong to different clades, suggesting that members of the same clade are involved in different functions. The highly specific expression of four *FvRALF* genes in male gamete and late stage of anther development (*FvRALF4, FvRALF5, FVRALF10, FvRALF1)* shown by the heatmap (Fig.2), suggests a role for these RALF genes in the ovule-pollen, cell-cell communication during the sequence of events precisely regulated during fertilization. A recent study reports that in Arabidopsis *At*RA*LF34* gene, expressed in the ovule, competes with *RALF4* and *RALF19*, expressed in the pollen tube, for binding to BUPs and ANXs receptors [13]. The interaction between AtRALF34, expressed by the female gamete, and receptor complex formed by BUPS1/2 and ANX1/2, present on pollen tube membrane leads to pollen tube rupture and sperm release [13]. The competitive binding of AtRALF4 and AtRALF19 to this receptor complex suggest that they have a redundant function in regulating pollen tube growth and integrity [16]. It is possible that these RALF genes functional redundancy is conserved in woodland strawberry.

Among the *FvRALF* genes expressed in flower and fruit organs *FvRALF1* and *FvRALF7* are the most highly expressed at the early stage of development in shoot apical meristem (SAM), floral meristem (FM) and receptacle meristem (RM), with *FvRALF1* also being the family member most highly expressed in the leaves. As for fruit, *FvRALF8* is the gene most highly expressed in mature fruits, whereas *FvRALF6, FvRALF7, FvRALF9* are transcribed during fruit growth and in the mature organ at 15 days post anthesis (15 DPA) both in Yellow Yonder and Red Rugen genotypes (the two *F. vesca* genotypes used for RNA seq), while *FvRALF3* and *FvRALF13* are more expressed in the mature fruits at 20 DPA. In particular *FvRALF1, FvRALF6, FvRALF7* and *FvRALF13* expression decreases during fruit development both in the inner and the outer tissues of fruit. On the contrary *FvRALF9* expression gradually increases with fruit growth.

In a recent work, Jia et al. [50] analyzed the expression of the woodland strawberry (*F. vesca*) Malectin Receptor Like Kinases (MRLK) also known as the *Catharanthus roseus* RLK-like proteins (CrRLK1Ls). *F. vesca* MRLKs are encoded by more than 60 genes, and more than 50% of these are expressed during fruit development. The majority of fruit *FvMRLK* genes are expressed at high level only at the early stage of fruit ripening, and decrease at ripe stages. In particular, Jia et al. [50], showed that transiently silencing and overexpression of *MRLK47* in strawberry fruit, severely affected ripening regulation. Moreover, a recent report describes how MRKL47 changes the sensitivity of ripening-related genes to ABA, a key hormone for strawberry fruit ripening [51]. Consistently, both RALF and ABA were found to be FERONIA-mediated cross-talk signals in stress-response and cell growth in *Arabidopsis* [52]. These data, together with an established role of FERONIA receptor in cell-wall integrity and Ca^2+^ signaling [53] both known to be important during fruit growth and ripening [54]-, and with the similar expression profile of FvMRLKs to the one observed here for *FvRALF1, FvRALF6, FvRALF7* and *FvRALF13,* support an important role for the RALF-MRLK signalling also for strawberry fruit development. Future studies should be conducted in order to demonstrate that RALF peptides and FvMRLK receptors interact *in vivo*.

### Expression profile of RALF genes in *Fragaria* x *ananassa* fruit and induction upon pathogen infection

RALF peptides are known to play a role in plant-pathogen interaction [28] since they were found to negatively regulate plant immunity response in *Arabidopsis* [12]. They were also found to be secreted by fungal pathogen as crucial virulence factors [26, 27]. Furthermore, it was reported t that genes homologous to *AtRALF33* were upregulated both in tomato (*Solanum lycopersicon)* and strawberry (*Fragaria x ananassa)* susceptible ripe fruits interacting with *Colletorichum gloeosporioides* and *C. acutatum,* respectively [30, 31, 33].

For this reason, the transcript levels of *FaRALF* genes were assessed in *Fragaria* x *ananassa* fruit at two different ripening stages and upon infection with two different fungal pathogens, *C. acutatum* or *B. cinerea.* Fruit *FaRALF* gene targets were chosen based on the *FvRALF* gene homologs expressed in fruit (Fig 2) since the *F.ve* progenitor is reported to have the most abundant expression level among the different subgenomes [36]. Therefore, primers were designed to amplify genes that are expressed in fruit (*FaRALF1-1, FaRALF3-1, FaRALF6-1, FaRALF7-1, FaRALF8-1, FaRALF9-1 and FaRALF13-1*), which included orthologs to *AtRALF33* (*FaRALF3-1, FaRALF8-1, FaRALF9-1* and *FaRALF13-1)*.

*FaRALF3* shows a significant expression increase induced by infection with both pathogens at the susceptible ripe stage, whereas *FaRALF9* expression decreased in white fruits upon *C. acutatum* but not upon *B. cinerea* infections (Fig 3). The expression of *FaRALF8* and *FaRALF13* were not affected by infection, since no significant difference between infected and control samples was observed either in white or red fruits (Fig 3). Out of the other *FaRALF* genes analyzed, only *FaRALF6* shows a clear downregulation in infected fruits at both ripening stage and pathogen species, while *FaRALF1 and FaRALF7* genes expression is significantly decreased only in red ripe stage with *B. cinerea* and in white stage with *C. acutatum,* respectively.

**Fig 3.**
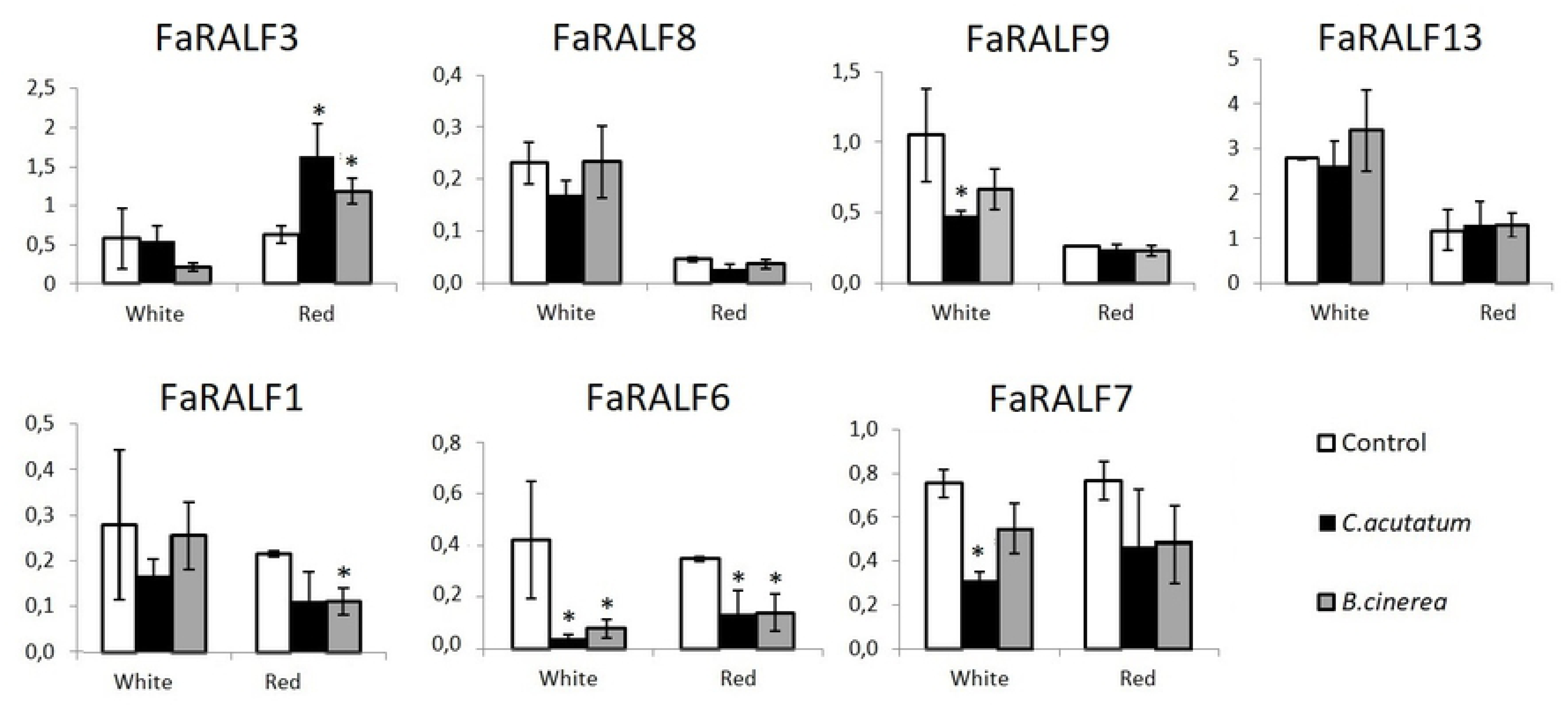
Expression of different RALF genes upon infection with pathogens. qRT-PCR analysis of different RALF genes (at the top the RALF-like 33 genes *FaRALF3, FaRALF9, FaRALF8* and *FaRALF13*) in Fxa strawberry fruits at different ripening stage (white and red) after 24 hours post infection with *C. acutatum* and *B. cinerea.* Histogram bars represent three biological replicates average and black lines represents standard deviations. T-student test between infected samples and control was used to calculate statistical significance. p< 0.05 (*).

The expression profiles of fruit RALF genes on *Fxa* strawberry fruits infected with two post-harvest pathogens are consistent with our previous results [31] showing that upon transient overexpression of a *FaRALF33-like* gene (here now named *FaRALF3)* the fruit susceptibility is affected. Notably, the *FaRALF3* and *FaRALF13* genes encode for mature peptides differing only for two aminoacids (S2 Figure) but respond differently to pathogen infection, suggesting a different role for these peptides.

Furthermore, the N-terminus sequence alignment of FaRALF1-1, FaRALF3-1, FaRALF7-1, FaRALF8-1a and FaRALF13 peptides with AtRALF23 (Fig 4A) shows that residues directly involved in AtRALF23 binding with AtLLG2 are conserved, suggesting that FaRALF peptides in strawberry fruits may also interact with the *Fxa* LLG2 homolog and *Fxa* FER Malectin Like Receptor Kinase (MRLK) to a heterotypic complex. Consistently, the homology models of FaRALF3-1 peptide interaction with FaMRLK47 - the MRLK mostly expressed in Fxa fruits [50] and with the *Fxa* LLG2 homolog (maker-Fvb3-4-snap-gene-34.65), shows that the structural components necessary to bind MRLK47 and LLG2 are conserved in FaRALF3 (Fig 4B and C), suggesting a similar binding mechanism and complex formation. In *Arabidopsis,* At*RALF23* binding to FERONIA and LLG proteins leads to negative immunity response regulation, however further analyses will be required to test FaRALF3-1 effectiveness binding to MRLK47 in fruits.

**Fig 4:**
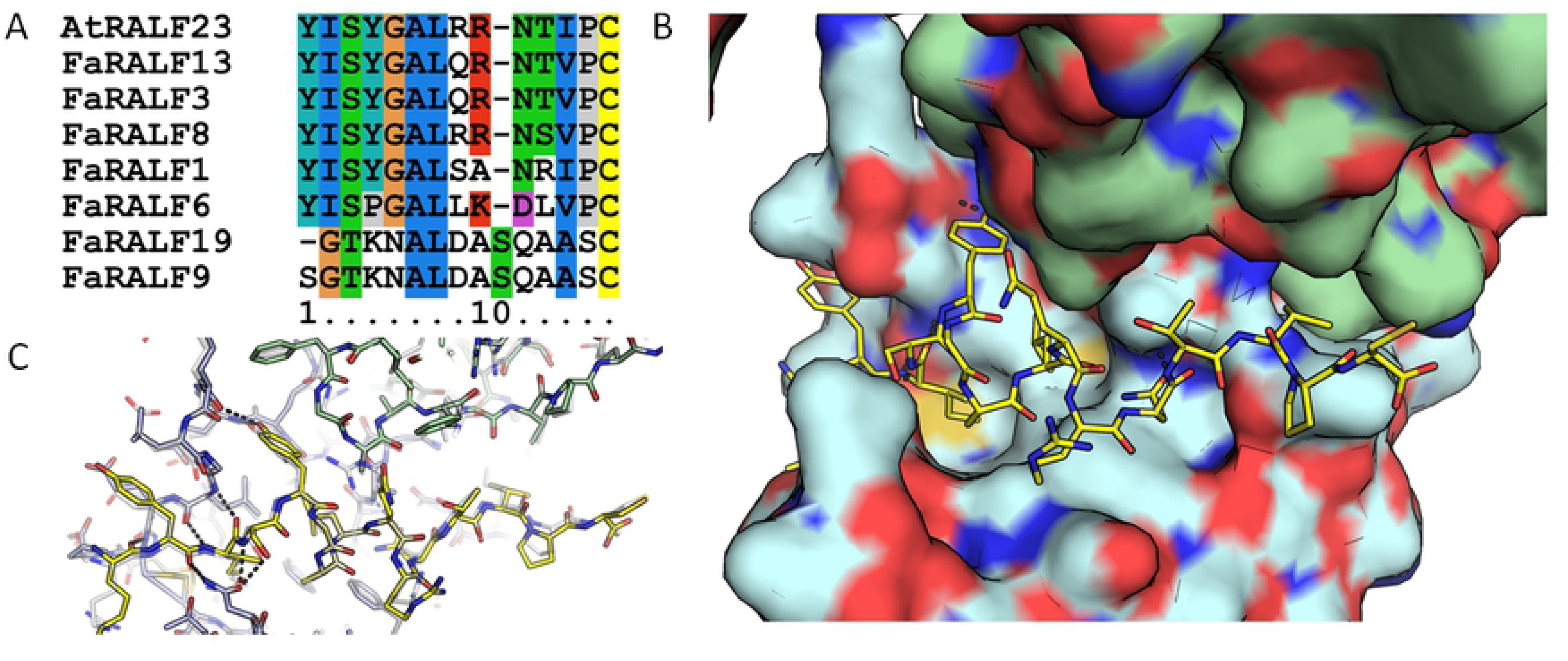
Model of FaRALF3, FaMRLK47 and LLG2. (A) multiple sequence alignment of FaRALFs and AtRALF23. (B) Model of the ternary complex of FaRALF3, FaMRLK47 and LLG2 based on At complex (PDB:6a5e). C. overlay of the Fa complex model and the At complex, showing high conservation of both the RALF and the two receptors.

Finally, the expression of *FaRALF1, FaRALF6* and *FaRALF9* are decreased during infection, thus it is possible that these *FaRALF* gene members may play different roles than *RALF3* homologs do during plant-pathogen interaction and immune response.

### *FaRALF3* promoter analysis in *Fragaria x ananassa* subgenomes and varieties

As shown above and reported in previous studies, RALF gene expression is triggered by different biotic and abiotic stimuli, however the signaling events regulating its expression are not yet known. In particular those responding to fungal pathogen infection. Identification of the promoter elements necessary for gene induction by fungal pathogens could provide insights on the transcription factors involved in immunity signaling and ultimately provide for the necessary knowledge to develop synthetic pathogen-responsive promoters to fight infections. Among *FaRALF* family genes, *FaRALF3* has shown clear upregulation in response to pathogen, and its overexpression in strawberry fruits is related to susceptibility [31]. For this reason *FaRALF3* was chosen for promoter characterization analysis.

To study *FaRALF3* putative promoter function in *Fragaria* x *ananassa*, the level of sequence conservation among the *Fxa* subgenomes of the region upstream RALF start codon was firstly assessed. *FaRALF3-1* sequence from the 3’UTR of upstream flanking gene (annotated as ‘*maker-Fvb2-2-augustus-gene-47.69-mRNA-1’*) and its ATG (590 bp) was used as input for BLASTn analysis against *Fragaria x ananassa cv.Camarosa* v1.0.a1. Five sequences were retrieved on *F. iinumae* Chr2-4, *F. nipponica* Chr2-1 and *F. viridis* Chr 2-3 (S4 Figure), indicating that the putative *FaRALF3* orthologs promoter sequences in the the *Fxa* subgenomes is highly conserved except for *F. viridis,* already reported to be the most divergent and silent in octoploid genome [36].

To study allelic variability, the *FaRALF3-1* putative promoter sequence similarity was assessed also in genomes of *Fragaria x ananassa* varieties with different susceptibility to fungal pathogens, the *cv. Florida Elyana* from Florida (U.S.A.) which is resistant to anthracnose disease [55], and *cv. Alba,* an italian variety which is highly susceptible to *C. acutatum* infection (https://plantgest.imagelinenetwork.com/it/varieta/frutticole/fragola/alba/59). The promoters were amplified with specific primers and five clones for each variety were sequenced and aligned with that of the v1.0.a1 genome sequence *cv. Camarosa* (S5 Figure). Only a single nucleotide polymorphism was detected between *cv. Alba* and Florida Elyana. This suggests that the function associated with the *Fa*RALF3-1 5’ upstream sequence in octoploid strawberry might be very important for expression regulation and also that the plant susceptibility to anthracnose disease cannot be associated with allelic polymorphisms in *FaRALF3-1* putative promoter.

### Prediction of *FvRALF3* promoter pathogen-responsive regulatory elements

The *FaRALF3-1* (from *F.vesca* subgenome) putative promoter sequence was chosen for pathogen-responsive regulatory element analysis because of the reported *F. vesca* subgenome dominance in octoploid genome [36]. Since *Fa*- and *FvRALF3* upstream putative regulatory sequences share 99% level of identity and the available *Fxa* pathogen-responsive transcriptome data have all been mapped onto *F. vesca* genome, the analysis of the *FaRALF3-1* promoter regulatory elements responsive to pathogen infection were carried out on *F. vesca* genome *FvRALF3* promoter.

For this, 656 bp of the F. vesca genomic sequence located between the stop codon of the *FvRALF3* upstream flanking gene on Chr2 and *FvRALF3* ATG starting codon, was analysed. *FvRALF3* putative promoter sequence was compared with known transcription factor binding sites of genes known to be regulated in *Fxa* strawberry fruit upon *C. acutatum* and *B. cinerea* infections [30, 45] in PLACE database [47] (Motif Scanning analysis). These latter sequences and RALF3 putative promoter were then scored for motif frequency (Motif Discovery analysis) (Fig 5).

**Fig 5.**
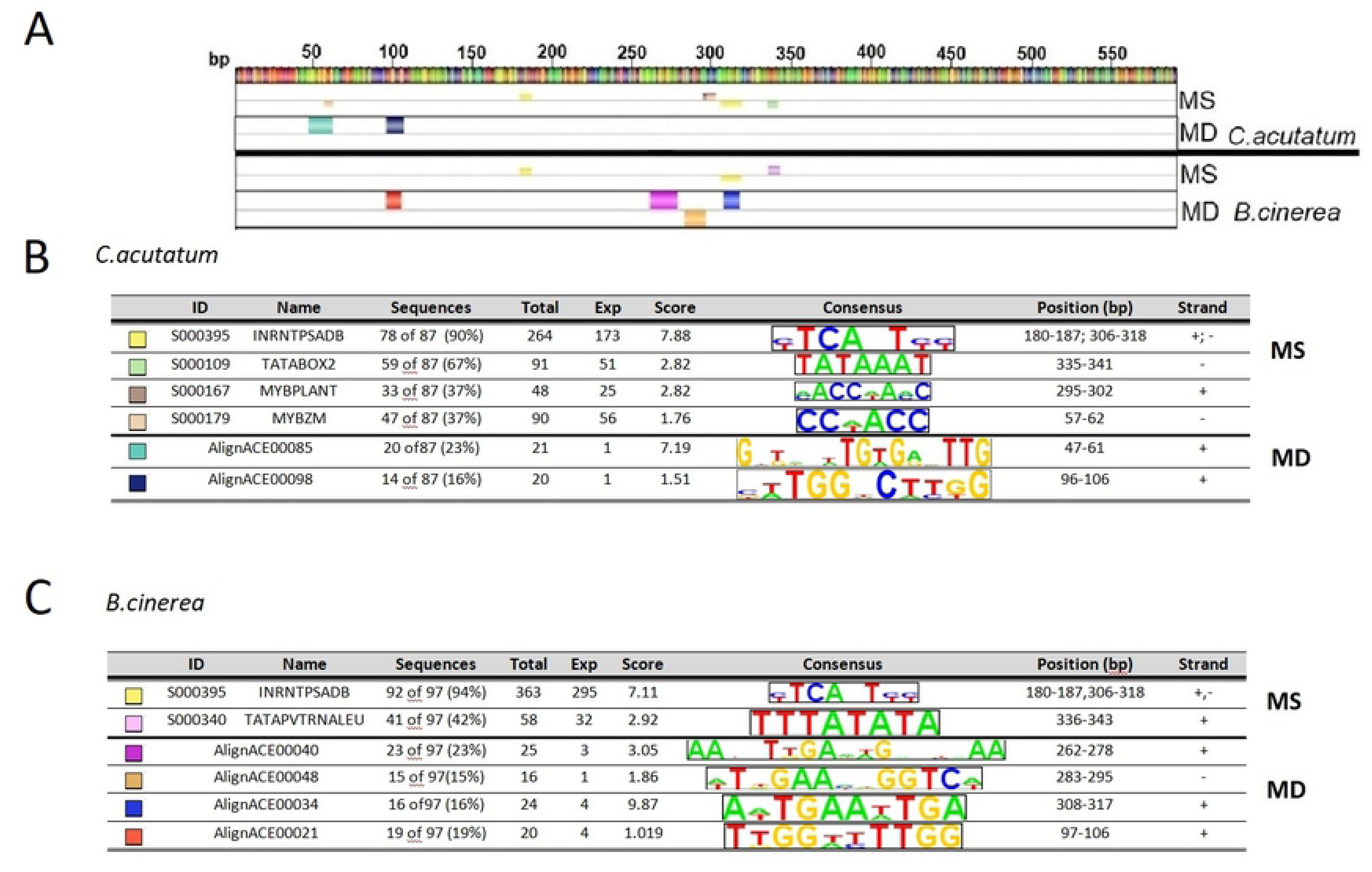
*In silico* analysis of predicted regulatory elements in pathogen-induced *Fragaria* genes. (A) From the top: FvRALF3 gene putative promoter, Motif Scanning (MS) and Motif Discovery (MD) outputs resulting from the analyses of *C. acutatum-*induced genes, and MS and MD outputs from the analysis of *B. cinerea-*induced genes after 24 hour post-infection. Small colored boxes in MS indicate known regulatory elements from PLACE database found to be significantly abundant in the group of sequences analyzed and in FvRALF3 putative promoter, while large coloured boxes in MD represent sequences found significantly enriched in the upstream sequences of genes analyzed. (B) and (C) are the color key respectively for *C. acutatum* and *B. cinereal* reported in (A). Codes and names of regulatory elements are reported, together with elements percentage abundance among sequences analyzed (Sequences), total count of elements found (total), element background frequency calculated as number of elements found in a third order random generated sequences (Exp), ranking score values calculated by MotifLab software using a binomial test with p-value threshold of 0.05 (Score). Consensus sequence of each element is shown along the position of the element in the putative FvRALF3 promoter (Position bp) and the strand in which it is found.

Motif scanning analysis of cis-acting elements mostly represented among fungal-induced genes revealed the presence of an element initially identified as initiator of PsaDb gene promoter (INRNTPSADB, PLACE ID: S000395), lacking TATA-box, which is common to almost all the sequenced analyzed and *FvRALF3* putative promoter (90% of the *C. acutatum* responsive genes promoter sequences and to 94% of *B. cinerea* ones) [56] (Fig. 5). Other TATA-like elements such as TATABOX2 (PLACE ID: S000109) and TATAPVTRNALEU (S000340), which have the role of recognition and initiator of transcription complex [57] were also found in several sequences (respectively in the 67% of *C. acutatum* and 42% of *B. cinerea* induced genes). Among the group of genes upregulated by *C. acutatum*, 33% and 47% have respectively a MYBPLANT (S000167) [58] and MYBZM (S000179) [59] regulatory elements, which are binding sites for MYB homolog. Additional analysis on the putative promoters of Arabidopsis *RALF33* (*AT4G15800*] and its homologous in tomato (*S. lycopersicum Solyc09g074890.1)*, revealed the presence of the same binding site (data not shown]. MYB proteins are a large family of transcription factor involved in regulation of many processes in plants, such as phenylpropanoid metabolism [60], ABA and JA signaling [61], responses to abiotic and biotic stress [62], cell death and circadian clock. MYB proteins generally interact with basic helix-loop helix (bHLH) family member and are regulated by cytosolic WD40 repeat proteins through formation of MYB/bHLH/WD40 dynamic complexes, which regulate various gene expression pathways [63]. Interestingly, MYB46 is involved in enhancing *B. cinerea* resistance through down-regulation of cell wall associated genes (CESA) during early stage infection [62]. Arabidopsis T-insertion mutants of genes regulated by MYB46, such as the a zinc-finger containing protein *gene zfp2,* the Basic Helix-Loop-Helix TF *bhlh99*, the AUX/IAA-type transcriptional repressor *pap2* and the *at1g66810 gene* coding for a Zing Finger Transcription Factor [TF], showed enhanced susceptibility to the necrotrophs fungal pathogens *B. cinerea* and *P. cucumerina* [34]. The transcriptional analysis of these four mutants revealed a coordinated upregulation of *RALF23, RALF24, RALF32* and *RALF33* [34] supporting the hypothesis of a MYB46 regulation of RALF gene expression.

A Motif Discovery analysis was performed using AlignACE algorithm [48] on the putative promoter sequences of both FvRALF3 and the identified *C. acutatum* and *B. cinerea* strawberry upregulated genes, and significantly overrepresented motifs were assessed. The motifs identified using MotifLab software are named ‘AlignACE’ followed by progressive numbers. For *C. acutatum* gene group the AlignACE00085 element (consensus GxTxxxTGTGAxTTG) was found in the 23% of sequences and in *FvRALF3* putative promoter, and is partially overlapping with MYBZM elements at the position between bases 57 and 62 of *FvRALF3* putative promoter. The AlignACE00098 (xxTGGxCTTGG) element was found in the 16% of *C. acutatum* upregulated genes and align to the elements AlignACE00021 (TTGGxxTTGG) found in *B. cinerea* upregulated group (19% of sequences analyzed). This suggests that in this position this element might be a regulatory component important for Fv*RALF3* fungal induced expression. Furthermore in *B. cinereal* gene group the AlignACE00040 (AAxxTTGAxxGxxxAA), AlignACE00048 (TxGAAxxGGTC)and AlignACE00034 (AxTGAAxTGA), located between bases 262 and 317 in *FvRALF3* putative promoter, were found significantly overrepresented.

### FvRALF3 promoter Agrobacterium-medieted reporter assay

In order to assess *FaRALF3* putative promoter function and identify possible pathogen-responsive regulatory elements, three progressive truncated fragments of *FaRALF3-1* upstream sequence consisting of the above described 588 bp sequence (T6), a 200 bp deletion (T4) and 400 bp deletion (T2) (Fig 6A), were cloned into pKGWFS7 vector and fused to two tandem reporter genes eGFP and β-glucoronidase (GUS) (S1 Figure). *Agrobacterium-*mediated transient transformation of white *Fxa* fruits was performed through injection of bacteria transformed with the three constructs. Fruits were infected with *C. acutatum* and analyzed for both reporter genes activity at 24 hpi, through quantification of eGFP expression in qRT-PCR and histochemical GUS assay for β-glucuronidase activity. GUS reporter activity, visualized as blue color of fruits, showed great variability among infected and mock-infected fruits (Fig 6B). Consistent to this, no significant difference was shown in the eGFP transcript level quantified in *C. acutatum* infected versus control fruit. This is possibly due to the fruit response to Agrobacterium itself, independently from the fungal pathogen. Indeed Agrobacterium can be perceived as a pathogen by the fruit and stimulate similar responses as *C. acutatum*, including those leading to FaRALF3 expression. The variability affecting strawberry *Agrobacterium*-mediated transformation depending on technical and environmental conditions has recently been described [64]. It was shown that the expression level of a reporter gene is normally distributed in an population of 30 treated fruits, with huge variation among different fruits. Other important factors affecting agroinfiltration methodology are the quantity of bacteria injected for each fruits, the stage of fruit ripening and the temperature and incubation time after transient transformation. In our experiments, maybe for all these reasons, it was not possible to infer the identity of *FaRALF3-1* promoter elements inducible by fungal pathogens such as *C. acutatum*, in agrobacterium-free systems. On the other hand, with respect to the different putative promoter sequence sizes tested, clearly the GUS activity and eGFP expression controlled by T4 and T6 sequences are measurable and comparable, whereas, under control of T2 element, both the GUS and eGFP become almost undetectable, suggesting that T4, comprising 400 bp sequence upstream *FaRALF3-1* ATG, contains the minimal promoter sequence elements necessary to drive reporter gene expression in strawberry fruits. According to Motif Scanning analysis, the T4 promoter fragment includes at least two regulatory elements known to be recognized by transcriptional activation complex, (TATA-boxes and Initiator of activation in TATA-less promoter) (Fig 6A and Fig 5). At the same time, the 200 bp sequence comprising the 3’ UTR of the Low PSII Accumulation (LPA1) and Tetratricopeptide (TRP) domains containing protein genes, are probably not determinant as expression regulatory sequences.

**Fig 6.**
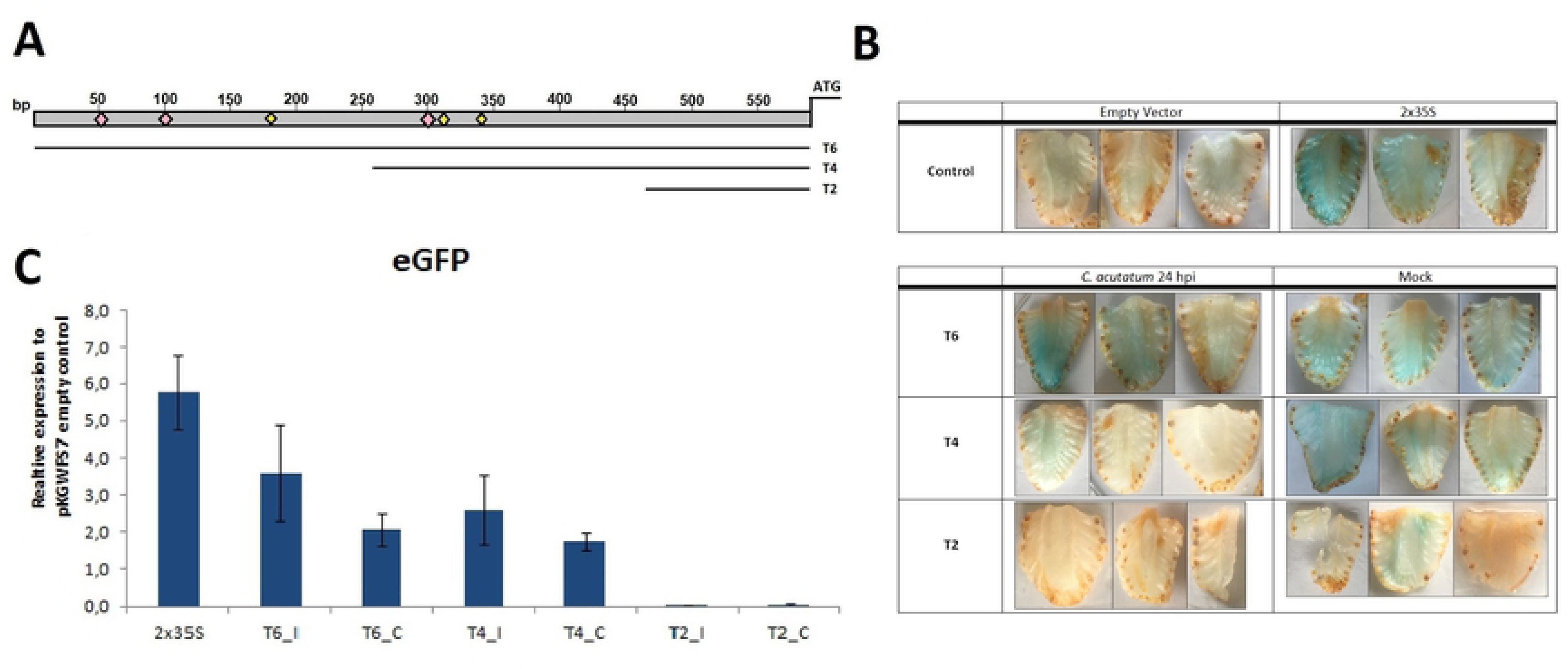
Dissection of *FaRALF3-1* promoter and *Agrobacterium*-mediated reporter assay. (A) Schematic representation of the promoter fragments used in this study. Pink squares indicate putative MYB-related regulatory elements, yellow squares indicates putative TATA-box related transcriptional activation elements. Histochemical GUS staining to detect β-glucoronidase activity in longitudinal fruit sections of Agroinfiltrated fruits. The table shows fruits transformed with negative (empty vector) and positive (double tandem p35S, 2×35S) controls. Fruits transformed with T6, T4 and T2 truncated promoter fragments were treated with *C. acutatum* or mock-treated and collected 24h post-infection. (B) Histochemical GUS staining of fruits to detect β-glucoronidase activity in longitudinal fruit sections of Agro-infiltrated fruits. The table shows fruits transformed with negative (empty vector) and positive (double tandem p35S, 2×35S) controls. Fruits transformed with T6, T4 and T2 truncated promoter fragments were treated with *C. acutatum* or mock-treated and collected 24h post-infection. Three replicates for each conitions are shown.(C) Histogram showing qRT-PCR quantitative analysis of eGFP reporter expression. Each bar represents the average of three biological replicates. Expression values were normalized to pKGWFS7 empty vector infilitarted fruits. (C) Mock-tretated samples, (I) *C. acutatum* infected

## CONCLUSIONS

Rapid Alkalinization Factors are small signal peptides with multiple roles in plant growth, fertilization and immunity regulations. RALF genes are upregulated in different plant hosts upon pathogens attack and RALF homologous genes are also expressed by many fungal pathogens as virulence factors, suggesting a role as susceptibility factors during plant pathogen interaction. The present work aimed to characterize the RALF gene family in *F. vesca* woodland and Fxa octoploid strawberries according to tissue specific expression and homology to *Arabidopsis* RALFs. The RALF gene family members distribution among *Fragaria x ananassa* subgenome is consistent with octoploid genome evolution, which is characterized by different TE activity in the different genome components and consequently different gene distribution. A putative involvement of MYB transcription factor as regulator of *FaRALF3-1* infection-inducibility is speculated based on *in silico* promoter characterization and MYB motif recognition. This is comprised in the 400 bp upstream the start codon of *FaRALF3-1* gene sequence. Since FaRALF3-1 presents the same conserved N-terminal sequence of AtRALF23 it is speculated that, consistently with motif conservation, FaRALF3-1 interaction with receptor FERONIA (MRLK47) and the coreceptor FaLLG2 follows the *Arabidopsis* complex interaction structure. Future research and methodology optimization will be required to identify specific pathogen-responsive elements.

## Aknowledgments

Protein studies was conducted at the University of Cambridge, UK, thanks to EMBO short fellowship number 7779. We thank Dr. Beata Blaszczyk and Aleksei Lulla for the kind help and support given during the period FN spent at University of Cambridge, UK.

We wish thank Prof. Bruno Mezzetti and his group lab members (Polytechnic University of Marche) for continuous support and suggestions on the experimental work carried on strawberry plants.

FN has been supported by PhD fellowship from MIUR (Ministry of Instruction, University and Research).

## Supporting Information

**S1 Figure. Map of pKGWFS7 plasmid used for promoter reporter assay.** Sm/Spr, spectinomycin resistance. Kan, kanamycin resistance. FvRALF3 promoter, T6,T4,T2, empty or 2×35S promoter according to experimental procedure. eFGPand GUS, chimera reporter gene formed by eGFP and β-glucoronidase ORFs in frame. T35S, terminator.

**S2 Figure. *F.vesca* RALF peptide sequences aligned in Clustal Omega.** Black boxes highlight conserved domains RRILA cleavage site, YISY activation site, and conserved cysteines (*).

**S3 Figure. *F.vesca* and *Fragaria x ananassa* RALF peptide sequences aligned in Clustal Omega.** Black boxes highlight conserved domains RRILA cleavage site, YISY activation site, and conserved cysteines (*).

**S4 Figure. Blastn output of *FaRALF3-1* upstream sequence against *Fragaria x ananassa cv. Camarosa* v1.0.a1 pseudomolecule using GDR.**

**S5 Figure. *FaRALF3-1* putative promoter sequence alignment in different *Fragaria x ananassa* variaties.** It was considered *Fragaria x ananassa cv. Florida Elyana from Florida (U.S.A)*, the Italian variety *cv.Alba* and the sequenced *cv. Camarosa (v1.0.a1)*.

**Table S1: List of *Fragaria x ananassa* RALF genes identified through *FvRALFs* BLASTx (GDR)**. In the table are listed for each gene, Chromosome localization ‘Chr’, subgenome localization ‘Subg.’, gene identification number used to name FaRALF in this work ‘geneID’, ‘*Fragaria x ananassa cv. Camarosa* transcript v1.0.a1 annotation’, ‘Genome location’ coordinates, classification in clades according to Campbell and Turner (2017) ‘clade’, *F. vesca* orthologous gene ‘Fv.Ort.’, ‘E-value’ and identity rate ‘identity’. ‘Gene ID’ column presents ‘*v1short name’* reporting gene v1 annotation abbreviation used to identify genes in Fig1A and ‘*orthology based’* nomenclature, assigned according to FvRALF orthology and chromosome lineage: −1, −2,−3,−4 after FvRALF orthologous name were used respectively to indicate *F. vesca, F. iinumae, F. nipponica* and *F. viridis* progenitors, progressive letters was used to name genes orthologous to the same FvRALF gene.

**TableS2.** List of primers used for RALFs qRT-PCR expression analysis In *Fragaria x ananassa* infected fruits, and for reporter gene expression in transient transformed fruits.

